# 5C-ID: Increased resolution Chromosome-Conformation-Capture-Carbon-Copy with in situ 3C and double alternating primer design

**DOI:** 10.1101/244285

**Authors:** Ji Hun Kim, Katelyn R. Titus, Wanfeng Gong, Jonathan A. Beagan, Zhendong Cao, Jennifer E. Phillips-Cremins

## Abstract

Mammalian genomes are folded in a hierarchy of compartments, topologically associating domains (TADs), subTADs and looping interactions. Currently, there is a great need to evaluate the link between chromatin topology and genome function across many biological conditions and genetic perturbations. Hi-C generates high quality, high resolution maps of looping interactions genome-wide, but is intractable for high-throughput screening of loops across conditions due to the requirement of an enormous number of reads (>6 Billion) per library. Here, we describe 5C-ID, an updated version of Chromosome-Conformation-Capture-Carbon-Copy (5C) with restriction digest and ligation performed in the nucleus (in situ Chromosome-Conformation-Capture (3C)) and ligation-mediated amplification performed with a new double alternating design. 5C-ID reduces spatial noise and enables higher resolution 3D genome folding maps than canonical 5C, allowing for a marked improvement in sensitivity and specificity of loop detection. 5C-ID enables the creation of high-resolution, high-coverage maps of chromatin loops in up to a 30 Megabase subset of the genome at a fraction of the cost of Hi-C.

## Introduction

Higher-order folding of chromatin in the 3D nucleus has been linked to genome function. Mammalian genomes are arranged in a nested hierarchy of territories [1], compartments [2–4], topologically associating domains [5–8] (TADs), subTADs [3, 9] and long-range looping interactions [10, 11]. Looping interactions have been linked to at least two mechanistically different modes of control over gene expression. First, enhancers can loop to distal target genes in a highly cell type-specific manner to facilitate their precise spatial-temporal regulation [12–15]. Second, long-range loops anchored by the architectural protein CTCF are often constitutive among cell types and form the structural basis for TADs/subTADs [9]. CTCF-mediated interactions connecting loop domains can create insulated neighborhoods that demarcate the search space of enhancers within the domain [16]. Specifically, CTCF anchored constitutive loops can prevent ectopic enhancer activation of genes outside of the domain or aberrant invasion of nonspecific enhancers into an inappropriate domain [16–20]. An active area of intense investigation includes the mapping of 3D loops genome-wide among hundreds of cell types, species and developmental lineages.

As genome-wide reference maps of looping interactions become widely available, a critical emerging goal will be to unravel the cause and effect relationship between looping and gene expression. Indeed, there is a great need in the field to build upon descriptive mapping studies and begin to perturb the 3D genome and evaluate the link of chromatin topology to function. One major limitation preventing progress is that genome-wide Hi-C technology requires more than six billion reads per replicate to obtain high quality, high resolution, genome-wide looping maps [3, 13, 21]. The financial and logistical difficulties of obtaining this read depth makes it intractable to conduct studies with a high number of samples with perturbations induced by genome editing or drug induction. Thus, there is a great need for a technology that creates ultra high-resolution 3D genome folding maps at a much lower cost.

Chromosome Conformation Capture Carbon Copy (5C) is proximity ligation technology pioneered by Dekker and colleagues [22, 23]. 5C adds a hybrid capture step to the classic Chromosome Conformation Capture (3C) method to facilitate the selection of all possible ligation products that occur only in a subset of the genome [22, 24–26]. Equivalent resolutions to Hi-C can be achieved at the fraction of the cost by only querying a subset of interactions in a 10-20 Megabase (Mb) subset of all genome-wide contacts [7, 9, 17, 27]. The ability to query a subset of genome contacts is important because genome-editing experiments are often conducted at only one specific location in the genome. The organizing principles governing genome folding can be queried at a key subset of loops without requiring the resources to map all loops genome wide, thus allowing many samples and perturbation conditions to be screened in a high-throughput manner.

Despite key advantages in the original 5C technique, it also has key challenges that have held back its widespread use, including: (1) the high number of cells (>40 million) required for the 3C template [2, 9, 27, 28], (2) the high amount of spatial noise caused by non-specific ligation products [29, 30] and (3) the non-comprehensive nature of the alternating primer design [7, 22–26, 31], resulting in many important interactions missing and a high degree of spatial noise. In the present study, we introduce two major updates to the 5C protocol that lead to a marked increase in resolution, decrease in spatial noise, and increased sensitivity and specificity of loop detection. We conduct a comparative analysis of in situ [3, 32] vs. canonical dilution 3C [2, 28] and a double alternating [17] vs. single alternating primer design [7, 22–26, 31] and report the downstream effect of these changes on 5C’s ability to detect bona fide looping interactions.

## Results

### Overview

A 5C experiment starts with preparation of the 3C template (**Figure 1A-B**). Chromatin is fixed within a population of cells with formaldehyde. In canonical dilution 3C [2, 28], cellular and nuclear membranes are disrupted and chromatin is digested in solution with a restriction enzyme (**Figure 1A**). Ligation is subsequently performed under dilute conditions that promote intra-molecular ligation. By contrast, *in situ* 3C [3, 32] involves restriction enzyme digest and ligation within intact nuclei. In both methods, cross-links are reversed and DNA is isolated to create the 3C template, which represents the genome-wide library of possible hybrid ligation junctions across a population cells (**Figure 1B**).

**Figure 1.**
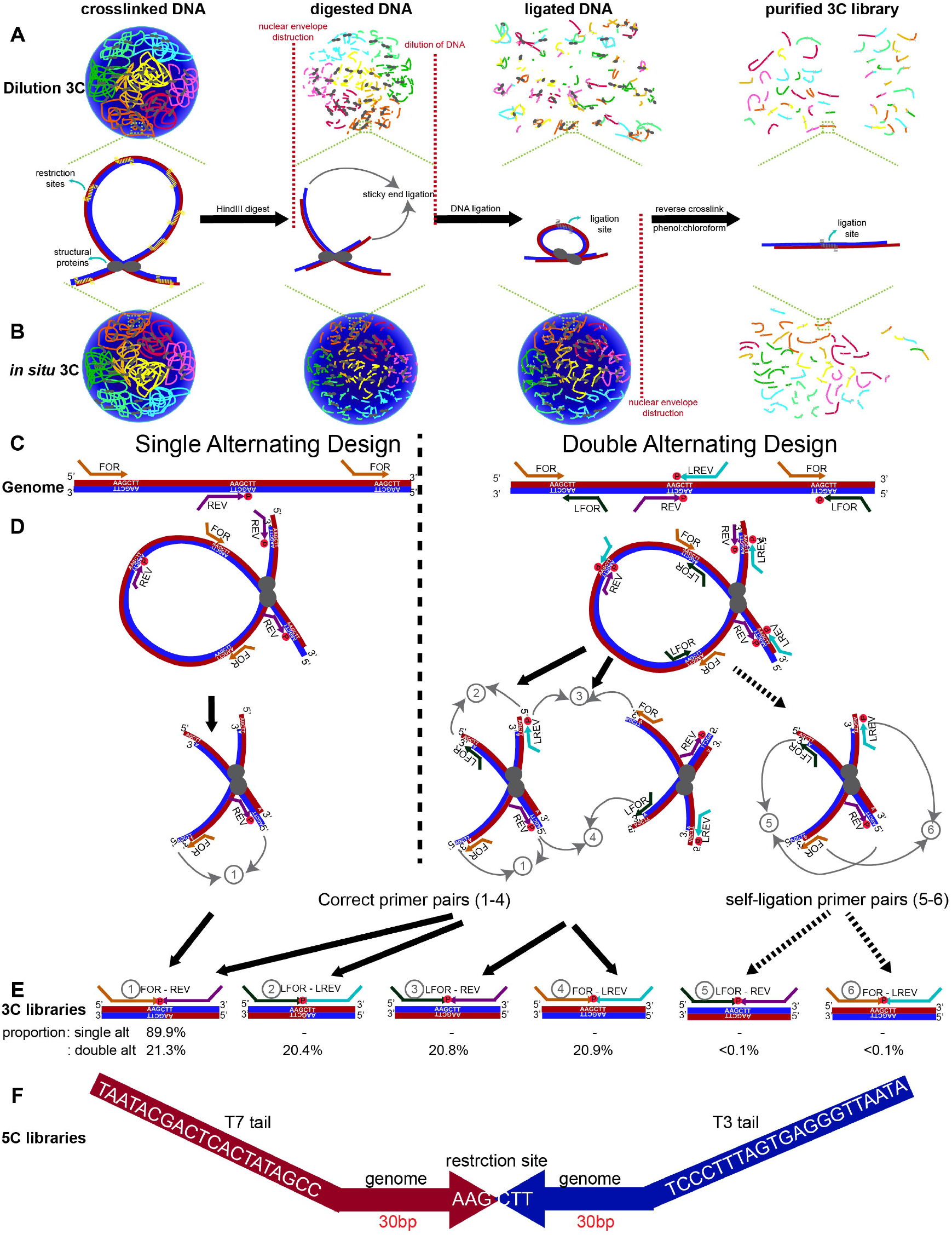
Overview of 5C-ID compared to canonical Chromosome-Conformation-Capture-Carbon-Copy (5C). (**A**) Schematic of dilution 3C. After chromatin fixation, the cellular and nuclear membranes are disrupted and chromatin is digested with a restriction enzyme in solution. After digestion, sticky ends are subsequently ligated under dilute conditions that promote intra-molecular ligation. (**B**) Schematic of *in situ* 3C. Unlike dilution 3C, chromatin digestion and ligation are performed in situ within intact nuclei. Chromatin colors illustrate DNA from independent chromosome territories. Vertical grey lines on DNA fragments illustrate ligation junctions. (**C**) Schematic illustrating single and double alternating 5C primer designs. Red and blue lines indicate sense and antisense DNA strands. (**D-E**) Schematic illustrating all possible correct 5C primer ligations from single and double alternating designs (1-4) as well as artifactual self-circle ligations between two ends of the same fragment (5-6). Percentages of each possible primer-primer pair orientation observed from a recent 5C experiment are provided. (**F**) Schematic illustrating the 5C amplicon from ligation-mediated amplification, including the T7/T3 universal tails, the 30 base pairs that uniquely bind to the genomic DNA and the religated Hindlll restriction sites.

The second half of the 5C protocol involves a hybrid capture step based on ligation-mediated amplification to select only a distinct subset of junctions from the genome-wide 3C library (**Figure 1C-F**). Canonical 5C [7, 22–26, 31] is built on an alternating primer design in which every other fragment is represented by either a Forward (FOR) primer binding to the sense strand or a Reverse (REV) primer binding to the antisense strand (**Figure 1C, left**). The single alternating design only queries approximately half of all ligation junctions in a target region because only FOR-REV primer ligation events are possible (**Figure 1D-E, left**). More recently, Dekker, Lajoie. and colleagues created a new double alternating primer design [17] which incorporates two additional "left-oriented’ primers, LFOR and LREV (**Figure 1C right**). The LFOR primer orientation is designed to the antisense strand on fragments also queried by REV primers, whereas the LREV primer orientation is designed to the sense strand on fragments also queried by FOR primers. Thus, the double alternating 5C primer design, there are now two primers representing each fragment, leading to 4 possible primer ligation orientations (FOR-REV, LFOR-LREV, LFOR-REV, FOR-LREV) and the query of nearly all fragment-fragment ligation events in an a priori selected Megabase (Mb)-scale genomic region (**Figure 1D-E right**).

### Double alternating primer design achieves increased loop detection sensitivity compared to single alternating design

We hypothesized that by using the double alternating design developed by Dekker and colleagues[17], we could markedly improve canonical 5C’s matrix resolution and loop detection sensitivity. To test this idea, we first started with a canonical dilution 3C template from pluripotent embryonic stem (ES) cells (detailed in **Materials and Methods**) and compared the quality of 5C libraries created at the same genomic region with both single alternating and double alternating primer designs. A tradeoff of the more comprehensive double alternating primer design is the possibility of artifactual ‘self-circles’ *(i.e.* ligation events between the 5’ and 3’ ends of the same restriction fragment; (5) and (6) in **Figure 1D-E right**). We counted the proportion of each possible primer ligation from the double alternating 5C experiment on a dilution 3C template from ES cells. There was an even distribution of ligation events across the four biologically informative primer-primer orientations ((1) FOR-REV: 21.3%, (2) LFOR-LREV: 20.4%, (3) LFOR-REV: 20.8%, (4) FOR-LREV: 20.9%). Importantly, self-circle ligation events ((5) LFOR-REV and (6) FOR-LREV from the same fragment) comprised only <0.1% of all primer ligations (**Figure 1E**), suggesting that the risk of self-ligation is very small.

We visually inspected 4 kb-binned heatmaps of 5C counts in Megabase-scale genomic regions around *Sox2* and *Zfp462* genes after matrix balancing and sequencing depth correction (detailed in **Materials and Methods**). We observed that the double alternating primer design results in marked improvement in specific, punctate looping signal between known long-range enhancer promoter-interactions compared to the single alternating primer design (**Figure 2A-B**). Double alternating 5C maps also showed less missing fragments than single alternating primer maps due to the increased complexity of ligation junctions that are queried and sequenced. In previous 5C studies, a smoothing window was required to reduce the blockiness of looping signal caused by missing ligation junctions [27, 33]. Here, with double alternating design, we use a 4 kb bin with no smoothing window and achieve punctate looping signal, little missing fragments and markedly reduced spatial noise (**Figure 2A-B**).

**Figure 2.**
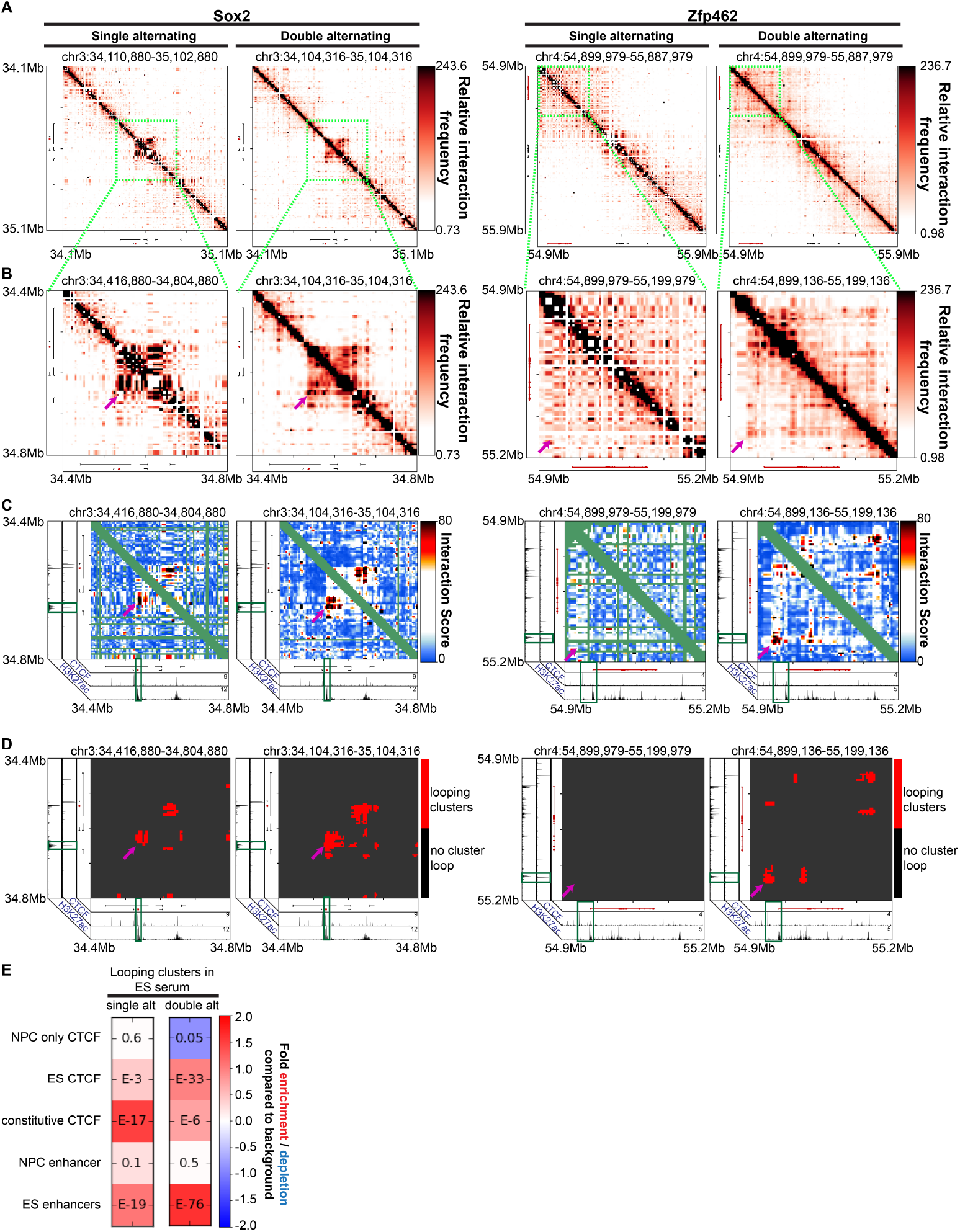
Improved 5C loop detection specificity with double vs. single alternating primer design. (**A**) Heatmaps binned at 4 kb matrix resolution showing relative chromatin interaction frequencies in 1 Mb regions surrounding *Sox2* and *Zfp462* genes across single alternating and double alternating primer designs in embryonic stem cells. Genes of interest are highlighted in red. (**B-D**) Zoomed-in heatmaps highlighting *Sox2* and *Zfp462* interactions with published pluripotency-specific enhancers. (**B**) Relative 5C interaction frequency after sequencing depth correction, binning and matrix balancing. **(C)** Interaction scores after distance-dependence and local background expectation correction and modeling. **(D)** Long-range looping interaction clusters after thresholding on interaction scores. ChIPseq tracks for CTCF and H3K27ac from embryonic stem cells are overlayed over the maps. **(E)** Enrichment of chromatin features at classified looping interactions relative to background interactions in 5C libraries using a single alternating primer design vs. double alternating primer design. P-values are calculated using Fisher’s exact test.

To further test our qualitative observation of increased looping sensitivity with the double alternating design, we also quantified chromatin looping interactions in each 5C dataset. We modeled binned interactions as a fold-enrichment relative to a background expected model based on distance dependence and local chromatin domain architecture (detailed in **Materials and Methods**). As previously published [27, 30, 33], we modeled these so-called Observed/Expected values with a parameterized logistic distribution and subsequently converted p-values to interaction scores (**Figure 2C**; detailed in **Materials and Methods**). After thresholding interaction scores, we clustered adjacent looping pixels into long-range looping interaction clusters (**Figure 2D**; detailed in **Materials and Methods**). Consistent with observations in **Figure 2A-B**, the interaction score and loop cluster maps also highlight punctate Sox2 and Zfp462 gene promoter-enhancer looping clusters (**Figure 2C-D**). As expected, the chromatin fragments anchoring the base of detected looping interactions contained high signal for H3K27ac, a chromatin modification known to demarcate active non-coding regulatory elements and active transcription start sites. Importantly, we identified key looping interactions between *Zfp462* and distal enhancers with the double alternating primer design that were not present with the single alternating design. The well-established Sox2-super enhancer interaction [5, 9, 27, 33–35] was detected by the single alternating design, but significantly more punctate and less blocky/noisy with the double alternating design.

Our loop detection analysis also suggests that the double alternating primer design would enable more precise discovery of epigenetic marks involved in looping. We intersected cell-type specific annotations of epigenetic marks from ES cells and primary neural progenitor cells (NPCs) [33] with our identified looping clusters (**Figure 2E**). Looping clusters in the double alternating 5C library are significantly enriched for ES-specific CTCF and ES-specific enhancers and depleted of NPC-specific CTCF. Consistent with the notion that looping specificity is improved with the double alternating design, we see that looping clusters in the single alternating 5C library are also enriched for ES-specific features but to a lesser degree than the double alternating design. Overall, these data indicate that the double alternating primer design allows for more sensitive detection of both strong and weak looping interactions.

### *In situ* 3C reduces spatial noise in 5C heatmaps compared to dilution 3C

We next assessed the quality of the double alternating 5C experiment using *in situ* 3C and dilution 3C templates. We prepared the *in situ* and dilution 3C templates from 2 million and 40 million ES cells cultured in 2i media, respectively, as previously reported (detailed in **Materials and Methods**). Both dilution and in situ 3C led to detection of previously reported looping interactions between *Sox2* and *Zfp462* and their target enhancers (**Figure 3A-D**, magenta arrowheads). Looping interactions from both *in situ* 3C and dilution 3C templates showed enrichment of ES-specific enhancers and ES-specific CTCF (**Figure 3E**). Although the enrichments and the number of loops appeared to be similar between the two templates, visual inspection of the maps revealed an extremely high degree of spatial noise and abnormal looping clusters from the dilution 3D template (**Figure 3A-D**). Spatial variance in dilution 3C was ~330x and ~280x higher than that of *in situ* 3C in the genomic regions around the *Sox2* and *Zfp462* genes, respectively (**Figure 3F-G**). *In situ* 3C resulted in a major improvement in spatial noise (**Figure 3F-G**) and led to looping interaction pixels grouped in more spherically shaped clusters with minimal background noise around the punctate looping pixels. These results indicate that *in situ* 3C is superior to dilution 3C in reducing overall spatial noise in heatmaps due to nonspecific ligation events.

**Figure 3.**
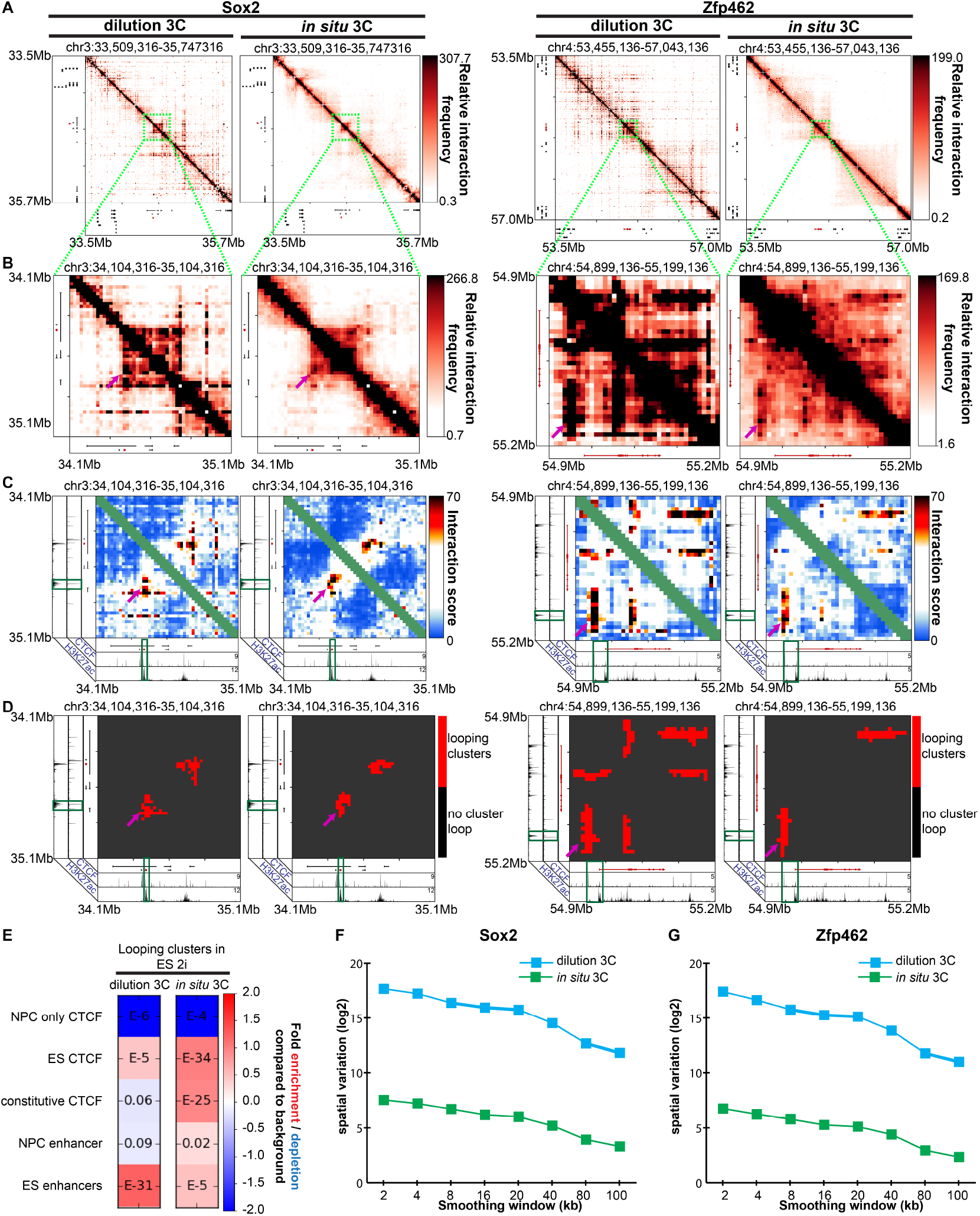
In situ 3C reduces spatial noise due to non-specific ligation products caused by dilution 3C. **(A)** Heatmaps binned at 8 kb matrix resolution showing relative chromatin interaction frequencies in 1 Mb regions surrounding *Sox2* and *Zfp462* genes. 5C libraries were created from *in situ* and dilution 3C templates from embryonic stem (ES) cells cultured in 2i media. Genes of interest are highlighted in red. (**B-D**) Zoomed-in heatmaps highlighting *Sox2* and *Zfp462* interactions with published pluripotency-specific enhancers. **(B)** Relative 5C interaction frequency after sequencing depth correction, binning and matrix balancing. **(C)** Interaction scores after distance-dependence and local background expectation correction and modeling. **(D)** Long-range looping interaction clusters after thresholding on interaction scores. ChIPseq tracks for CTCF and H3K27ac from embryonic stem cells are overlayed over maps. **(E)** Enrichment of chromatin features at classified looping interactions relative to background interactions in 5C libraries using a single alternating primer design vs. double alternating primer design. P-values are calculated using Fisher’s exact test. (**F-G**) Spatial variance of binned contact matrices around **(F)** *Sox2* and **(G)** *Zfp462* genes as a function of smoothing window size.

### Combined implementation of a double alternating primer design and *in situ* 3C allows for the use of lower genome copies than canonical 5C

We observed that implementing a double alternating primer design and *in situ* 3C noticeably improves the quality and resolution of our 5C heatmaps by reducing background noise and allowing for more sensitive detection of chromatin looping interactions (**Figure 2-3**). Therefore, we hypothesized that we could lower the genome copies required for loop detection by combining the two improvements. The advantage of lowering the required number of genome copies is that lower cell number 5C could be performed in the future, opening up opportunities for conducting 5C analysis on rare cell types and human tissue samples.

We performed double alternating 5C on an *in situ* 3C template made from ES cells cultured in 2i media. Canonical 5C typically performs the ligation-mediated amplification step on 200,000 genome copies (~590 ng) of the mouse 3C template. We tried 590ng, 245 ng, 120 ng, 12 ng, and 2.5 ng of the same *in situ* 3C library prepared from mouse ES cells in 2i media, representing 200,000, 100,000, 50,000, 5,000 and 1,000 mouse genome copies, respectively. To ensure that the total DNA mass did not affect 5C primer binding and ligation efficiencies, we mixed 3C templates with an excess of salmon sperm DNA (to a total DNA mass of 1,500 ng).

Visual inspection of heatmaps revealed that the 3C template mass could be reduced to 50,000 genome copies and still sensitively detect all gold-standard looping interactions (**Figure 4A-D**, Supplementary **Figure 1A-D**). Notably, the quality of chromatin looping signal is drastically reduced when the number of genome copies is further lowered to 5,000 and 1,000 (**Figure 4A-D**, **Supplementary Figure 1A-D**). Consistent with this result, quantitative chromatin enrichments were similar for 50,000-200,000 genome copies, but did not show interpretable results at 1,000-5,000 genome copies (**Figure 4E**). Since there were not any looping clusters called in the 5C libraries using 5,000 and 1,000 genome copies (**Figure 4E, G**), no enrichment or depletion was detected. Spatial noise was generally comparable with 200,000-50,000 genome copies, but was notably higher in libraries prepared with 1,000 and 5,000 genome copies (**Figure 4H**). Altogether, these data demonstrate that simultaneous implementation of a double alternating primer design and *in situ* 3C allows for successful 5C using lower genome copies. The implication of these results is that 5C might be performed on smaller cell populations in future studies.

**Figure 4.**
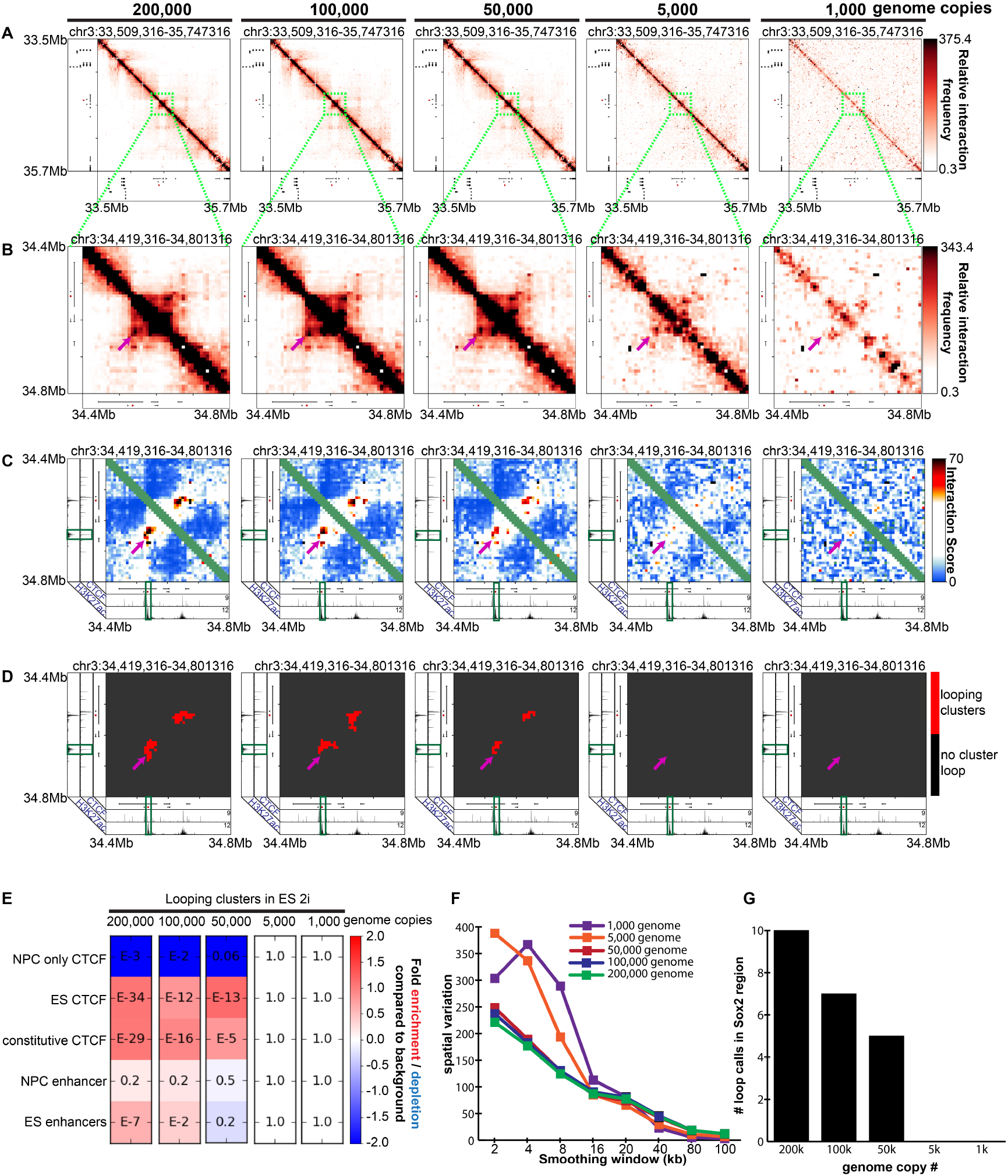
Combined usage of an in situ 3C template and double alternating primer design lowers the number of 3C template genome copies required to produce high quality 5C data. **(A)** Heatmaps binned at 8 kb matrix resolution showing relative chromatin interaction frequencies in 1 Mb regions surrounding the *Sox2* gene. 5C libraries were created from a double alternating design and *in situ* 3C templates (200,000, 100,000, 50,000, 5,000, 1,000 genome copies) from embryonic stem (ES) cells cultured in 2i media. Genes of interest are highlighted in red. (**B-D**) Zoomed-in heatmaps highlighting *Sox2* interactions with published pluripotency-specific enhancers. **(B)** Relative 5C interaction frequency after sequencing depth correction, binning and matrix balancing. **(C)** Interaction scores after distance-dependence and local background expectation correction and modeling. **(D)** Long-range looping interaction clusters after thresholding on interaction scores. ChIPseq tracks for CTCF and H3K27ac from embryonic stem cells are overlayed over maps. **(E)** Enrichment of chromatin features at classified looping interactions relative to background interactions in 5C libraries using a single alternating primer design vs. double alternating primer design. P-values are calculated using Fisher’s exact test. **(F)** Spatial variance of binned contact matrices at various genome copies as a function of smoothing window size. **(G)** Looping cluster numbers in the Sox2 region as a function of genome copy number.

## Discussion/Conclusions

The invention of the canonical 5C procedure by Dekker, Dostie and colleagues enabled the creation of high-resolution, high-coverage 3D genome folding maps from a subset of the genome (up to ~30 Mb) at a fraction of the cost of Hi-C [7, 9, 17, 27]. Due to the markedly reduced cost, 5C is poised to have high utility in addressing the significant unmet need of comprehensive inquiry of the folding of all fragments within a large genomic locus across hundreds of biological perturbation conditions. The ability to create loop resolution maps across thousands of gene editing perturbations is essential for testing the functional relationship between genome structure and function.

Despite these key advantages, canonical 5C has been limited in looping detection sensitivity and specificity due to the alternating primer design which only queries half the ligation junctions and non-specific ligations leading to a low signal to noise ratio. In the present study, we present an updated version of the classic 5C procedure, 5C in situ double alternating (5C-ID). 5C-ID implements a double alternating primer design [17] and *in situ* 3C [3, 32], resulting in markedly increased sensitivity for looping signal detection and reduced off-target non-specific ligation junctions. Double alternating primers comprehensively bind to all possible ligation junctions [17] missed by the single alternating design [7, 22–26, 31], leading to markedly improved loop detection sensitivity. Moreover, by conducting restriction digestion and ligation steps of 3C *in situ* in the nucleus, we dramatically reduced spatial noise cause by known nonspecific ligations from classic dilution 3C[2, 3, 28–30, 32]. By combining these two changes, we were also able to maintain loop detection sensitivity with reduced genome copies. While canonical 5C was performed on 40-100 million cells[9, 15, 36], only 2 million cells were used here for 5C-ID. The evidence that genome copies can be decreased to 50,000, which suggests pellets of ~25,000-50,000 cells or possibly less are possible in the future for high quality 5C maps. Thus, 5C-ID creates high-coverage, high-quality heatmaps at loop resolution at a fraction of the cost and opens the future potential for low cell number analysis from rare cell types and human tissues.

## Acknowledgements

We thank Job Dekker and Bryan Lajoie for providing us access to the My5C software and assistance in designing double alternating 5C primers. J. Phillips-Cremins is a New York Stem Cell Foundation (NYSCF) Robertson Investigator and an Alfred P. Sloan Foundation Fellow. This work was funded by The New York Stem Cell Foundation (J. Phillips-Cremins), the Alfred P. Sloan Foundation (J. Phillips-Cremins), the NIH Director’s New Innovator Award (1DP2MH11024701; J. Phillips-Cremins), a 4D Nucleome Common Fund grant (1U01HL12999801; J. Phillips-Cremins) and a joint NSF-NIGMS grant to support research at the interface of the biological and mathematical sciences (1562665; J. Phillips-Cremins).

## Authors’ Contributions

JEPC and JHK conceived of the study. JHK, WG and JAB performed 5C. KRT implemented the computational pipeline. JEPC, JHK, KRT wrote the manuscript

## References

[1] T. Cremer, C. Cremer, Chromosome territories, nuclear architecture and gene regulation in mammalian cells, Nature Reviews Genetics 2 (2001) 292.

[2] E. Lieberman-Aiden, N.L. van Berkum, L. Williams, M. Imakaev, T. Ragoczy, A. Telling, I. Amit, B.R. Lajoie, P.J. Sabo, M.O. Dorschner, R. Sandstrom, B. Bernstein, M.A. Bender, M. Groudine, A. Gnirke, J. Stamatoyannopoulos, L.A. Mirny, E.S. Lander, J. Dekker, Comprehensive Mapping of Long-Range Interactions Reveals Folding Principles of the Human Genome, Science 326(5950) (2009) 289–293.

[3] Suhas S.P. Rao, Miriam H. Huntley, Neva C. Durand, Elena K. Stamenova, Ivan D. Bochkov, James T. Robinson, Adrian L. Sanborn, I. Machol, Arina D. Omer, Eric S. Lander, Erez L. Aiden, A 3D Map of the Human Genome at Kilobase Resolution Reveals Principles of Chromatin Looping, Cell 159(7) (2014) 1665–1680.

[4] J. Fraser, C. Ferrai, A.M. Chiariello, M. Schueler, T. Rito, G. Laudanno, M. Barbieri, B.L. Moore, D.C. Kraemer, S. Aitken, S.Q. Xie, K.J. Morris, M. Itoh, H. Kawaji, I. Jaeger, Y. Hayashizaki, P. Carninci, A.R. Forrest, C.A. Semple, J. Dostie, A. Pombo, M. Nicodemi, Hierarchical folding and reorganization of chromosomes are linked to transcriptional changes in cellular differentiation, Molecular Systems Biology 11(12) (2015).

[5] J.R. Dixon, S. Selvaraj, F. Yue, A. Kim, Y. Li, Y. Shen, M. Hu, J.S. Liu, B. Ren, Topological domains in mammalian genomes identified by analysis of chromatin interactions, Nature 485(7398) (2012) 376–380.

[6] C. Hou, L. Li, Zhaohui S. Qin, Victor G. Corces, Gene Density, Transcription, and Insulators Contribute to the Partition of the *Drosophila* Genome into Physical Domains, Molecular Cell 48(3) (2012) 471–484.

[7] E.P. Nora, B.R. Lajoie, E.G. Schulz, L. Giorgetti, I. Okamoto, N. Servant, T. Piolot, N.L. van Berkum, J. Meisig, J. Sedat, J. Gribnau, E. Barillot, N. Blüthgen, J. Dekker, E. Heard, Spatial partitioning of the regulatory landscape of the X-inactivation centre, Nature 485 (2012) 381.

[8] T. Sexton, E. Yaffe, E. Kenigsberg, F. Bantignies, B. Leblanc, M. Hoichman, H. Parrinello, A. Tanay, G. Cavalli, Three-Dimensional Folding and Functional Organization Principles of the *Drosophila* Genome, Cell 148(3) (2012) 458–472.

[9] Jennifer E. Phillips-Cremins, Michael E.G. Sauria, A. Sanyal, Tatiana I. Gerasimova, Bryan R. Lajoie, Joshua S.K. Bell, C.-T. Ong, Tracy A. Hookway, C. Guo, Y. Sun, Michael J. Bland, W. Wagstaff, S. Dalton, Todd C. McDevitt, R. Sen, J. Dekker, J. Taylor, Victor G. Corces, Architectural Protein Subclasses Shape 3D Organization of Genomes during Lineage Commitment, Cell 153(6) (2013) 1281–1295.

[10] S. Chambeyron, W.A. Bickmore, Chromatin decondensation and nuclear reorganization of the HoxB locus upon induction of transcription, Genes & Development 18(10) (2004) 1119–1130.

[11] A.L. Sanborn, S.S.P. Rao, S.-C. Huang, N.C. Durand, M.H. Huntley, A.I. Jewett, I.D. Bochkov, D. Chinnappan, A. Cutkosky, J. Li, K.P. Geeting, A. Gnirke, A. Melnikov, D. McKenna, E.K. Stamenova, E.S. Lander, E.L. Aiden, Chromatin extrusion explains key features of loop and domain formation in wild-type and engineered genomes, Proceedings of the National Academy of Sciences 112(47) (2015) E6456–E6465.

[12] M.H. Kagey, J.J. Newman, S. Bilodeau, Y. Zhan, D.A. Orlando, N.L. van Berkum, C.C. Ebmeier, J. Goossens, P.B. Rahl, S.S. Levine, D.J. Taatjes, J. Dekker, R.A. Young, Mediator and cohesin connect gene expression and chromatin architecture, Nature 467(7314) (2010) 430–435.

[13] F. Jin, Y. Li, J.R. Dixon, S. Selvaraj, Z. Ye, A.Y. Lee, C.A. Yen, A.D. Schmitt, C.A. Espinoza, B. Ren, A high-resolution map of the three-dimensional chromatin interactome in human cells, Nature 503(7475) (2013) 290–4.

[14] B.D. Pope, T. Ryba, V. Dileep, F. Yue, W. Wu, O. Denas, D.L. Vera, Y. Wang, R.S. Hansen, T.K. Canfield, R.E. Thurman, Y. Cheng, G. Gulsoy, J.H. Dennis, M.P. Snyder, J.A. Stamatoyannopoulos, J. Taylor, R.C. Hardison, T. Kahveci, B. Ren, D.M. Gilbert, Topologically associating domains are stable units of replication-timing regulation, Nature 515(7527) (2014) 402–405.

[15] A. Sanyal, B.R. Lajoie, G. Jain, J. Dekker, The long-range interaction landscape of gene promoters, Nature 489 (2012) 109.

[16] Jill M. Dowen, Zi P. Fan, D. Hnisz, G. Ren, Brian J. Abraham, Lyndon N. Zhang, Abraham S. Weintraub, J. Schuijers, Tong I. Lee, K. Zhao, Richard A. Young, Control of Cell Identity Genes Occurs in Insulated Neighborhoods in Mammalian Chromosomes, Cell 159(2) (2014) 374–387.

[17] D. Hnisz, A.S. Weintraub, D.S. Day, A.-L. Valton, R.O. Bak, C.H. Li, J. Goldmann, B.R. Lajoie, Z.P. Fan, A.A. Sigova, J. Reddy, D. Borges-Rivera, T.I. Lee, R. Jaenisch, M.H. Porteus, J. Dekker, R.A. Young, Activation of proto-oncogenes by disruption of chromosome neighborhoods, Science 351(6280) (2016) 1454–1458.

[18] Y. Guo, Q. Xu, D. Canzio, J. Shou, J. Li, David U. Gorkin, I. Jung, H. Wu, Y. Zhai, Y. Tang, Y. Lu, Y. Wu, Z. Jia, W. Li, Michael Q. Zhang, B. Ren, Adrian R. Krainer, T. Maniatis, Q. Wu, CRISPR Inversion of CTCF Sites Alters Genome Topology and Enhancer/Promoter Function, Cell 162(4) (2015) 900–910.

[19] Darío G. Lupiáñez, K. Kraft, V. Heinrich, P. Krawitz, F. Brancati, E. Klopocki, D. Horn, H. Kayserili, John M. Opitz, R. Laxova, F. Santos-Simarro, B. Gilbert-Dussardier, L. Wittler, M. Borschiwer, Stefan A. Haas, M. Osterwalder, M. Franke, B. Timmermann, J. Hecht, M. Spielmann, A. Visel, S. Mundlos, Disruptions of Topological Chromatin Domains Cause Pathogenic Rewiring of Gene-Enhancer Interactions, Cell 161(5) (2015) 1012–1025.

[20] W.A. Flavahan, Y. Drier, B.B. Liau, S.M. Gillespie, A.S. Venteicher, A.O. Stemmer-Rachamimov, M.L. Suvà, B.E. Bernstein, Insulator dysfunction and oncogene activation in IDH mutant gliomas, Nature 529 (2015) 110.

[21] J.H.I. Haarhuis, R.H. van der Weide, V.A. Blomen, J.O. Yáñez-Cuna, M. Amendola, M.S. van Ruiten, P.H.L. Krijger, H. Teunissen, R.H. Medema, B. van Steensel, T.R. Brummelkamp, E. de Wit, B.D. Rowland, The Cohesin Release Factor WAPL Restricts Chromatin Loop Extension, Cell 169(4) (2017) 693–707.e14.

[22] J. Dostie, T.A. Richmond, R.A. Arnaout, R.R. Selzer, W.L. Lee, T.A. Honan, E.D. Rubio, A. Krumm, J. Lamb, C. Nusbaum, R.D. Green, J. Dekker, Chromosome Conformation Capture Carbon Copy (5C): A massively parallel solution for mapping interactions between genomic elements, Genome Research 16(10) (2006) 1299–1309.

[23] J. Dostie, Y. Zhan, J. Dekker, Chromosome Conformation Capture Carbon Copy Technology, Current Protocols in Molecular Biology, John Wiley & Sons, Inc. 2001.

[24] J. Dostie, J. Dekker, Mapping networks of physical interactions between genomic elements using 5C technology, Nature Protocols 2 (2007) 988.

[25] M.A. Ferraiuolo, A. Sanyal, N. Naumova, J. Dekker, J. Dostie, From cells to chromatin: Capturing snapshots of genome organization with 5C technology, Methods 58(3) (2012) 255–267.

[26] N.L. van Berkum, J. Dekker, Determining Spatial Chromatin Organization of Large Genomic Regions Using 5C Technology, in: P. Collas (Ed.), Chromatin Immunoprecipitation Assays: Methods and Protocols, Humana Press, Totowa, NJ, 2009, pp. 189–213.

[27] Jonathan A. Beagan, Thomas G. Gilgenast, J. Kim, Z. Plona, Heidi K. Norton, G. Hu, Sarah C. Hsu, Emily J. Shields, X. Lyu, E. Apostolou, K. Hochedlinger, Victor G. Corces, J. Dekker, Jennifer E. Phillips-Cremins, Local Genome Topology Can Exhibit an Incompletely Rewired 3D-Folding State during Somatic Cell Reprogramming, Cell Stem Cell 18(5) (2016) 611–624.

[28] J. Dekker, K. Rippe, M. Dekker, N. Kleckner, Capturing Chromosome Conformation, Science 295(5558) (2002) 1306–1311.

[29] E. Yaffe, A. Tanay, Probabilistic modeling of Hi-C contact maps eliminates systematic biases to characterize global chromosomal architecture, Nature Genetics 43 (2011) 1059.

[30] T.G. Gilgenast, J.E. Phillips-Cremins, Systematic evaluation of statistical methods for identifying looping interactions in 5C data, bioRxiv (2017).

[31] B.R. Lajoie, N.L. van Berkum, A. Sanyal, J. Dekker, My5C: web tools for chromosome conformation capture studies, Nature Methods 6 (2009) 690.

[32] T. Nagano, Y. Lubling, T.J. Stevens, S. Schoenfelder, E. Yaffe, W. Dean, E.D. Laue, A. Tanay, P. Fraser, Single-cell Hi-C reveals cell-to-cell variability in chromosome structure, Nature 502(7469) (2013) 59–64.

[33] J.A. Beagan, M.T. Duong, K.R. Titus, L. Zhou, Z. Cao, J. Ma, C.V. Lachanski, D.R. Gillis, J.E. Phillips-Cremins, YY1 and CTCF orchestrate a 3D chromatin looping switch during early neural lineage commitment, Genome Research (2017).

[34] Y. Li, C.M. Rivera, H. Ishii, F. Jin, S. Selvaraj, A.Y. Lee, J.R. Dixon, B. Ren, CRISPR Reveals a Distal Super-Enhancer Required for Sox2 Expression in Mouse Embryonic Stem Cells, PLoS ONE 9(12) (2014) e114485.

[35] B. Bonev, N. Mendelson Cohen, Q. Szabo, L. Fritsch, G.L. Papadopoulos, Y. Lubling, X. Xu, X. Lv, J.-P. Hugnot, A. Tanay, G. Cavalli, Multiscale 3D Genome Rewiring during Mouse Neural Development, Cell 171(3) (2017) 557–572.e24.

[36] Emily M. Smith, Bryan R. Lajoie, G. Jain, J. Dekker, Invariant TAD Boundaries Constrain Cell-Type-Specific Looping Interactions between Promoters and Distal Elements around the *CFTR* Locus, The American Journal of Human Genetics 98(1) 185–201.

[37] X. Wu, D.A. Scott, A.J. Kriz, A.C. Chiu, P.D. Hsu, D.B. Dadon, A.W. Cheng, A.E. Trevino, S. Konermann, S. Chen, R. Jaenisch, F. Zhang, P.A. Sharp, Genome-wide binding of the CRISPR endonuclease Cas9 in mammalian cells, Nat Biotech 32(7) (2014) 670–676.

[38] M. Imakaev, G. Fudenberg, R.P. McCord, N. Naumova, A. Goloborodko, B.R. Lajoie, J. Dekker, L.A. Mirny, Iterative correction of Hi-C data reveals hallmarks of chromosome organization, Nature Methods 9 (2012) 999.

